# Somatic structural variant formation is guided by and influences genome architecture

**DOI:** 10.1101/2021.05.18.444682

**Authors:** Nikos Sidiropoulos, Balca R. Mardin, F. Germán Rodríguez-González, Shilpa Garg, Adrian M. Stütz, Jan O. Korbel, Erez Lieberman Aiden, Joachim Weischenfeldt

## Abstract

The occurrence and formation of genomic structural variants (SV) is known to be influenced by the 3D chromatin architecture, but the extent and magnitude has been challenging to study. Here, we apply Hi-C to study chromatin organization before and after induction of chromothripsis in human cells. We use Hi-C to manually assemble the derivative chromosomes following the massive complex rearrangements, which allowed us to study the sources of SV formation and their consequences on gene regulation. We observe an *action-reaction* interplay whereby the 3D chromatin architecture directly impacts on the location and formation of SVs. In turn, the SVs reshape the chromatin organization to alter the local topologies, replication timing and gene regulation in *cis*. We show that genomic compartments and replication timing are important determinants for juxtaposing distant loci to form SVs across 30 different cancer types with a pronounced abundance of SVs between early replicating regions in uterine cancer. We find that SVs frequently occur at 3D loop-anchors, cause compartment switching and changes in replication timing, and that this is a major source of SV-mediated effects on nearby gene expression changes.

## Introduction

Understanding the mechanistic forces that shape somatic aberrations in cancer genomes can provide important basic and clinically relevant information on the causes of the disease. Massive genome sequencing efforts have profiled recurrent patterns of somatic mutations from tumor biopsies, though they can only capture a snapshot of the series of somatic aberrations that typically accumulate over several years or even decades. Mutations are shaped by stochastic, mechanistic and selective forces, and the sources of single-nucleotide variants (SNVs) have been explored extensively (Lawrence et al. 2014; Polak et al. 2015; Gonzalez-Perez et al. 2019; Pich et al. 2018). These studies have shown a strong association between SNV occurrence and one-dimensional features of the genome, including local chromatin, replication timing and transcription factor binding.

In contrast to SNVs, SVs can impact multiple genes through gene loss, disruption, gene-gene fusion formation, amplification, or dysregulation of the cis-regulatory landscape, in some cases through reorganising topology-associated domains (TADs), which can lead to enhancer hijacking (Weischenfeldt et al. 2017, 2013; Northcott et al. 2014). A pan-cancer study found relatively few TAD-affecting SVs to be associated with changes in nearby gene expression (Akdemir et al. 2020), which is in line with a general realisation that changes in gene expression does not necessarily depend on changes in contact domains (Despang et al. 2019; Ghavi-Helm et al. 2019). The sources of SVs in cancer have also proved challenging to disentangle, partly due to their complex nature and orders-of-magnitude lower occurrence frequency compared to SNVs (Drier et al. 2013; Yang et al. 2013; Wala et al. 2017; Rheinbay et al. 2020). SVs are formed through double-strand breaks (DSBs) and illegitimate ligation of normally distant loci in the genome or following replication-mediated mechanisms. SVs can be simple (e.g. deletions, duplications, translocations) or highly complex, as in the case of chromothripsis(Stephens et al. 2011; Rausch et al. 2012a), which leads to chromosome shattering and reordering of genomic segments from kilobases in size to whole chromosomes.

The 3D nuclear organization is known to be important in gene regulation through bringing genomically distant loci in close, physical proximity (Nora et al. 2012; Le Dily et al. 2014). Several lines of evidence have suggested that SV occurrence is linked with chromatin architecture (Lupiáñez et al. 2015; Canela et al. 2017; Gothe et al. 2019). Early cancer studies using fluorescence in situ hybridization (FISH) found that the proximity in 3D genome space influences the probability of forming fusion gene products (Roix et al. 2003; Lukásová et al. 1997). On a more global scale, the effects of genome architecture on SVs in cancer genomes have been inferred from somatic copy number aberrations (Fudenberg et al. 2011; De and Michor 2011) and WGS (Akdemir et al. 2020; Harewood et al. 2017; Dixon et al. 2018) using reference maps of genome organization (Fudenberg et al. 2011; De and Michor 2011). Moreover, SVs can alter the 3D organization, which can have direct consequences on gene regulation (Weischenfeldt et al. 2017; Northcott et al. 2017; Lupiáñez et al. 2015; Affer et al. 2014; Akdemir et al. 2020; Dixon et al. 2018). However, it remails unsolved to what extent prior 3D contacts in the same cells impact on somatic SVs formation in human cells, and secondly, how these acquired SVs impact on gene expression in cis. This had to date represented a challenge due to the lack of model systems to study the relationship between genome organization before and after SV occurrence in the same population of cells. Additionally, the lack of information on the physical composition of the chromosomes in the cell after multiple rearrangements have occurred, presents an obstacle. To appreciate the consequences of SVs on genome architecture and gene regulation, it is therefore desirable to construct a ‘derivative chromosome’ that represents the order in which genomic regions have been stitched together following complex SV formation. To shed light on the interplay between SVs and genome organization, we used Hi-C to compare and assemble human cells before and after chromothripsis, which enabled us to resolve complex SVs in an allele-specific manner from Hi-C (Lieberman-Aiden et al. 2009). We leverage this to investigate key properties of the 3D genome folding conformation, how it impacts SV formation across 2,700 cancer genomes and, ultimately, how SVs feed-back on the 3D genome architecture to alter gene and genome regulation properties.

## Results

### Reconstructing the derivative chromosomes from highly rearranged genomes

Doxorubicin (dox), a widely used chemotherapy medication to treat cancer, traps Topoisomerase II (TOP2) on DNA, causing TOP2-linked double-strand breaks (DSBs) which can result in the formation of massive, genome-wide SVs (Tewey et al. 1984; Fornari et al. 1994). We have previously developed a model system that uses dox treatment to induce chromothripsis in human cells, characterised by massive genomic rearrangements in a single cell cycle (Mardin et al. 2015). Our model system use TP53-deficient retinal pigment epithelial cell line RPE-1, a near diploid cell line widely used in genomic instability studies (Passerini et al. 2016; Santaguida et al. 2017; Wilhelm et al. 2019; Drainas et al. 2020). We performed mate-pair whole-genome sequencing (WGS) and Hi-C sequencing on the wild-type (C93), two TP53-deficient (C29 and DCB2; maternal) and on four dox-treated (BM175, BM178, BM838 and BM780, daughter) clones showing signs of massive rearrangements (Figure 1a and Figure S1a, Supp. Table S1-2). Hi-C maps can be leveraged to provide information on SVs (Harewood et al. 2017; Dixon et al. 2018; Jacobson et al. 2019) since physical proximity mediated by SVs are reflected in novel contacts in the Hi-C (Harewood et al. 2017; Dixon et al. 2018; Jacobson et al. 2019) map (Figure 1). We used Hi-C analysis to identify 306 SVs, the majority of which were also identified by mate-pair sequencing (Figure S2b-d). These SVs were explicitly occurring in the daughter cells, many of which were clustered on a few chromosomes (Figure S1a, Supp. Table S3-4), a typical feature of chromothripsis. Key criteria for chromothripsis are the prevalence of SVs on a single haplotype, minimal positional recurrence of breakpoints and random orientation of SVs (Korbel and Campbell 2013). To this end, we used our Hi-C data to phase the SVs (Tourdot and Zhang 2019), which confirmed that all SVs were associated with the same haplotype (Figure S1b-c), further supporting that the SVs are induced in a single cell cycle and represent a bona fide example of chromothripsis (Korbel and Campbell 2013). As expected, our analysis of four different daughter cells revealed no breakpoint recurrence and random orientation of SVs (Fig S2c), supporting that these events are associated with a more general source of genomic instability and not disruption of specific genomic sites or selection of specific SVs. Mutations in key DNA repair enzymes can give rise to specific patterns of complex SVs in cancer genomes (Li et al. 2020; ICGC/TCGA Pan-Cancer Analysis of Whole Genomes Consortium 2020). We next tested whether specific types of complex SVs were detectable in the genomes besides chromothripsis. We recently developed computational approaches to classify complex SVs into different categories including chromoplexy, templated insertions and breakage-fusion bridge cycle, as well as chromothripsis (Li et al. 2020; Cortés-Ciriano et al. 2020). Applying the methodologies to our highly rearranged samples only revealed chromothripsis as the mechanism of formation, further supporting a random breakpoint formation process and that no other faulty DNA repair-based processes are active in these genomes.

**Figure 1:**
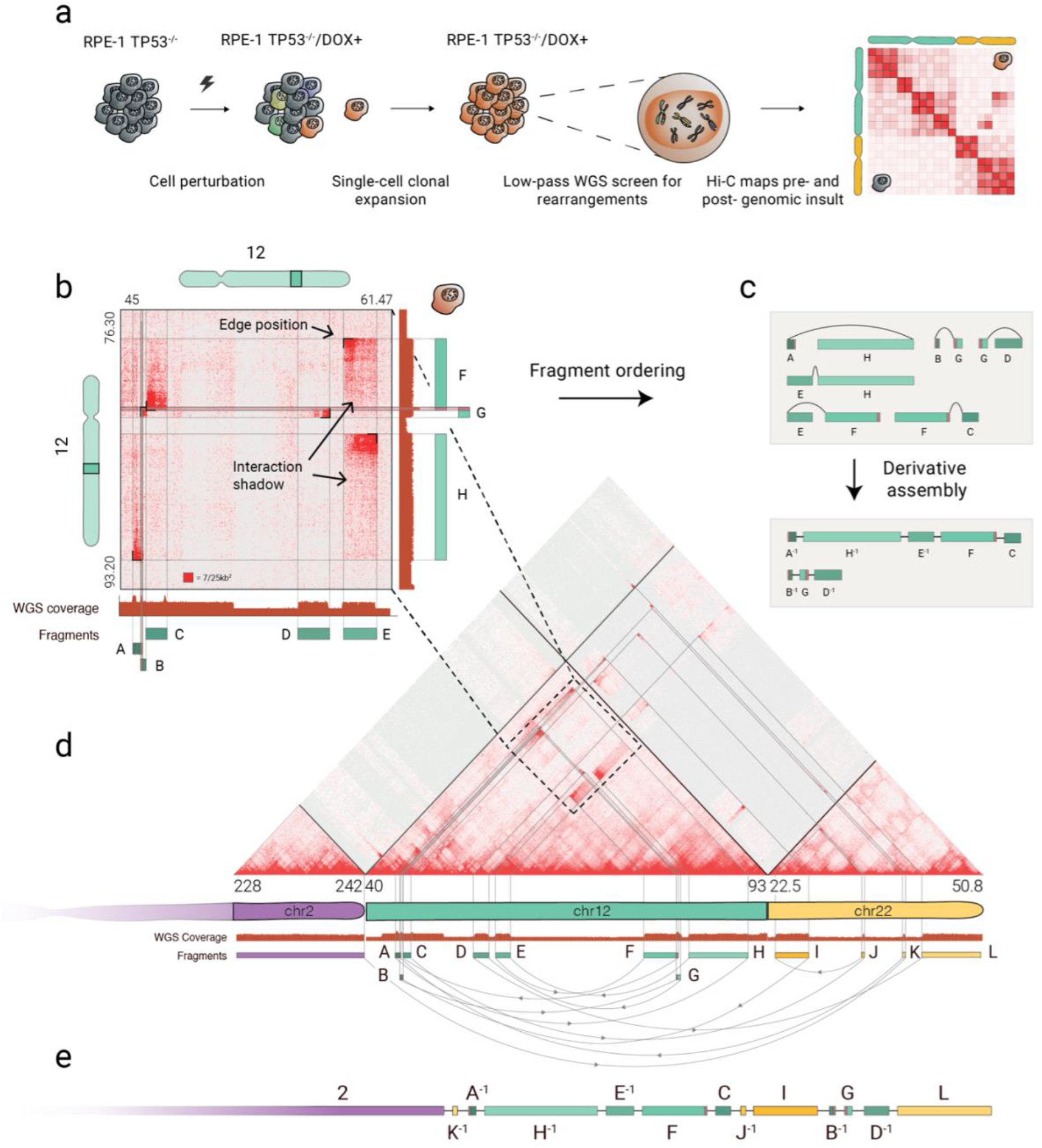
Hi-C-based assembly of a highly rearranged human cell line model system. a) Schematic overview of CAST (Complex Alterations after Selection and Transformation) coupled with in situ Hi-C. Briefly, *TP53*-deficient RPE-1 cells are perturbed with doxorubicin (DOX), followed by single-cell sorting and expansion. Surviving populations are screened for genomic rearrangements with low-depth WGS. Hi-C libraries from control (maternal) and treated (daughter) clones are sequenced to identify acquired and drug-induced rearrangements. b) Annotation of fragment size and edge positions. The latter is annotated at peaks of ectopic contacts between two distal sites. Tracing the interaction shadow that originates at the interaction peak reveals the length of the rearranged fragment. Duplicated regions, highlighted at fragments F and G, exhibit distinct interaction patterns and produce non-orthogonal ectopic interactions, as revealed by the interaction profile between fragments C-F and B-G respectively. The genomic regions depicted on the Hi-C maps correspond to the highlighted regions on the chromosome ideograms. c) (upper) Following the above described annotation strategy, we create a graph of nodes (fragments) and edges (rearrangements) for the ectopic interactions of panel (b). (lower) Traversing the graph produces the derivative assembly. −1 denotes fragment inversion. Duplicated regions that are shared between fragments A-B and F-G are colored in pink. d) The Hi-C map depicts rearrangements within and across chromosomes 2, 12 and 22. Straight diagonal lines on the Hi-C map trace the position of the SV calls back to the reference genome. Following the graph method in b-d, we annotated 12 rearranged DNA fragments (A-L) and their juxtapositions, represented as green and yellow rectangles, and directed arches respectively. We omitted five small segments of chromosome 12 for visualization purposes. See the complete derivative chromosome in Figure S2h. e) Using chromosome 2 as a starting point, we walk along the annotated edges and segments to produce the assembly of the derivative 2-12-22 chromosome.

Having established that the SVs in our samples were random and compatible with chromothripsis, we next sought to reconstruct the derivative chromosomes as they appeared in the genomes of the rearranged cells. This is challenging, due to the highly rearranged nature and heterozygous state of the genomes, and no automated methodologies are currently able to reconstruct the derivative chromosomes from such genomes. To this end, we used our Hi-C data to perform reference-guided manual assembly on the chromothriptic daughter cells to obtain the derivative chromosomes. Hi-C is particularly tractable due to the long-range genomic contact information that can be utilized to order genomic fragments and assign them to whole derivative chromosomes (Dudchenko et al. 2017; Harewood et al. 2017; Ghavi-Helm et al. 2019). The long-range information can assist in connecting even repeat-rich sequences, which are impossible to assemble with standard short-read paired-end sequencing.

Two key sources of information encoded in the ectopic Hi-C contact maps formed the main guidelines of our manual assembly. First, as it has been previously described (Harewood et al. 2017; Dixon et al. 2018), the breakpoint location of SVs can be identified as peaks of high-intensity ectopic interactions, which we term the edge position (Figure 1b). Second, and most importantly, the length and type of SV can be identified from the decay in contact frequency originating at the breakpoint site, which we term interaction shadow. The orientation of the interaction shadow can be used to identify the particular SV type, such as deletion, duplication and inversion-type (Figure 1b and Figure S2a). We initially focused on the highly rearranged chromosomes 12 and 22 of one of the daughter clones, BM178, to map all juxtapositions of the rearranged fragments in the derivative chromosome. We found that the interaction shadow is also helping to resolve conflicts across overlapping signals, as seen in the highlighted region between fragments F and G in Figure 1b. Fragment F has enriched contacts with fragment C, observed as high intensity regions on the Hi-C map, and depleted with B, while the opposite is true for G. Only the highlighted region has contact enrichment across all four fragments. This can only occur if the region was duplicated with a copy existing in both fragments F and G, each one with its own distinct interaction signature. The predicted duplication is confirmed by a copy-number gain in the WGS coverage track. A similar scenario is observed between fragments A and B, and their interactions with fragments G and H.

We used these two features, Hi-C edge position and interaction shadows, to create an assembly graph composed of genomic segments, represented as nodes, and SVs represented as edges. Traversing the graph produces the derivative chromosome (Figure 1c). We identified 13 genomic segments of chromosome 12q stitched in seemingly random orientations with four segments from chromosome 22q (Figure 1de and Figure S2h). The SVs and segment annotations are in alignment with copy-number switches and loss-of-heterozygosity identified by WGS (Figure 1b and Figure S3d). Moreover, we found evidence that the derivative chromosome resides at the telomere of chromosome 2 (Figure S2e), supported by contact frequency enrichment between fragments from chromosomes 12 and 22, and the q arm of chromosome 2 (Figure S2f). To validate the assembly, we used Juicebox Assembly Tools (JBAT) (Dudchenko et al. 2018) on allele-specific Hi-C maps to reconstruct a part of the predicted derivative chromosome, which confirmed the formation of ectopic domains between fragments E, F and H (Figure S2g).

The derivative chromosome assemblies allowed us to gain insight into the complex constituents of a highly rearranged, allele-specific cancer genome. We found that the derivative 2-12-22 chromosome involved the q-arm of chromosome 22 and fragments of 12q. The chromosome 22p arm was ligated to the remaining q arm of 12, while 12p acquired the q-arm of the X-chromosome (Figure S3a-b). We additionally identified another highly rearranged derivative 10-15-7 chromosome as well as 7-15, 11-13, 17-19 and 10-X (Figure S3c). The 10-X fusion was the only large-scale rearrangement inherited from WT RPE-1 cells (Janssen et al. 2011).

### Sites of structural variants are linked with pre-existing genomic contacts

Having established the physical composition of the derivative chromosome, we next turned to the 3D genome organization prior to chromothripsis. Since we generated WGS and Hi-C data from the maternal (before chromothripsis) and derivative cell lines (after chromothripsis), we went on to examine the impact on genomic contacts in 3D and SV formation. WGS on the derivative cell lines was performed on an early passage after chromothripsis induction, whereas Hi-C was performed approximately 20 passages later. This difference in passage timing between the library preparation of WGS and Hi-C, allowed us to dissect SVs that occurred as a direct result of dox treatment (induced) or due to subsequent genomic instability (spontaneous). Dox-induced SVs are expected to be clonal, present in all cells and evident in both WGS and Hi-C assays while spontaneous SVs are expected to be more subclonal and may only be supported by Hi-C. We detected 22 spontaneous events (Supp. Table S5) on two daughter clones, 13 on chromosome 10 of BM175 (Figure S4a) and 9 on chromosomes 11, 12 and 18 of BM178 (Figure S4b) that were not associated with copy-number switches on WGS data. We also detected 14 pre-existing SVs in C29 which were excluded from the spontaneous SV analysis. We computed the contact frequency in a 25×25kb window around induced and spontaneous rearrangements to create a clonality metric for all SVs. This clonality analysis revealed significantly lower contact frequencies for spontaneous SVs, characterizing them as subclonal events (Figure S4c). The two groups of SVs, induced and spontaneous, gave us a unique opportunity to identify the mechanistic basis for SVs associated with inhibition of TOP2 and SVs associated with subsequent genomic instability in the daughter cell genomes (Figure S4d). We used these data to ask whether and to what extent linear and 3D features of the genome were associated with juxtaposition of distal genomic regions and formation of SVs.

The genome can be partitioned into A and B compartments, associated with open and closed chromatin, respectively (Lieberman-Aiden et al. 2009); (Dixon et al. 2015). We utilized public available Hi-C data from 8 different cell lines (Rao et al. 2014) with different cell-of-origin to compute tissue-specific and conserved compartment domains (Figure 2a). In total, 95% of all compartments were conserved across different tissues (see Methods for details). Consistently, we found that the majority of SVs occur in conserved compartments, irrespective of whether they were dox-induced or spontaneous (Figure 2b). Interestingly, whereas the TOP2-linked SVs were more likely to occur in A-compartments (p=3.5e-13, Fisher’s exact test), the spontaneous SVs showed no preferential compartment-occurrence (Figure 2c, left).

**Figure 2:**
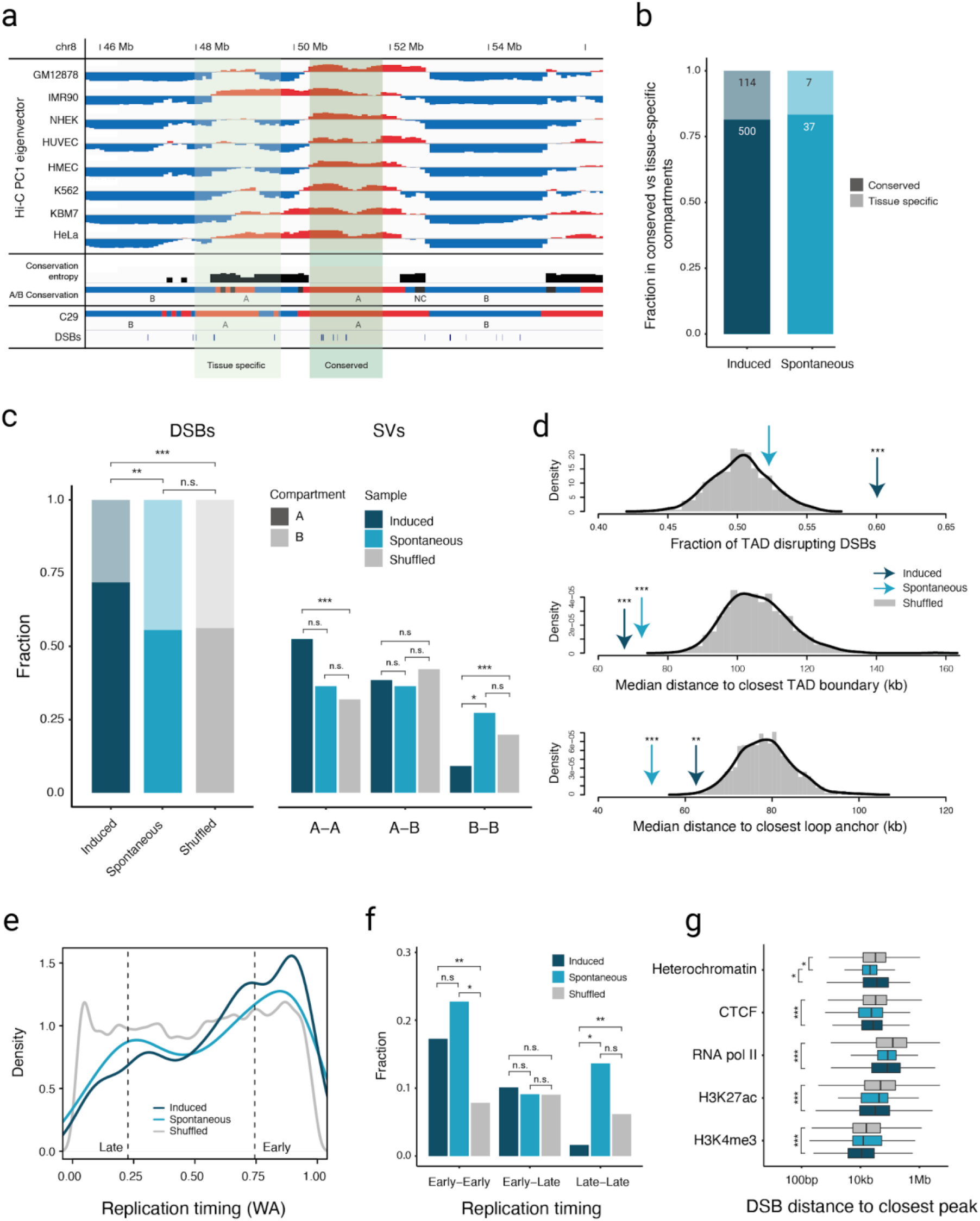
Genome architecture features at SV sites. a) DSBs associated with conserved and tissue-specific compartments. PC1 eigenvectors from 8 human cell lines (Rao et al. 2014), with signs adjusted so that positive values correspond to A compartment. A/B conservation track annotates every 100kb bin to a compartment it was found to be in the same state in >4 samples. Compartment entropy reflects the compartment concordance across the cell lines, with higher values representing less conserved compartments. DSBs in tissue-specific compartments overlap non-conserved regions or regions where the C29 compartment annotation disagrees with the A/B conservation track. Red and blue colors correspond to A and B compartments, respectively. NC = Not Conserved. b) Distribution of DSBs in conserved and tissue-specific compartments between induced and spontaneous rearrangements. 114 out of 614 (18.6%) and 7 out of 37 (18.9%) occurred in tissue-specific compartments. c) (left) Induced DSBs (*n* = 610) are highly enriched within A compartments compared to the spontaneous (*n* = 44) and shuffled (*n* = 588,388) set. ** p-value < 0.01, *** p-value < 0.001, Fisher’s exact. (right) SVs are enriched in A to A and depleted in B to B in the induced SV set compared to the shuffled, while spontaneous SVs are evenly distributed across A and B compartments. * p-value < 0.05, *** p-value < 0.001, post-hoc Fisher’s exact, FDR corrected. d) Observed and expected distribution of DSBs across features that reflect genome organization. (top) Induced DSBs are enriched inside TADs (*n* = 6,165) while spontaneous are enriched in inter-domain loci. Both of them occur significantly closer to insulating factors, such as TAD boundaries (middle, *n* = 12,326) and chromatin loop anchors (bottom, *n =* 17,374) than the shuffled sets. * p-value < 0.05, ** p-value < 0.01, *** p-value < 0.001 e) Replication timing Weighted Average (WA) values at DSBs show enrichment of induced (dark blue, *n* = 609) and spontaneous (light blue, *n* = 44) breaks during early replication compared to the permuted set (grey, *n* = 586,044). f) SVs occur significantly more frequently between Early to Early RT regions for both induced and spontaneous rearrangements. In contrast, Late to Late RT SVs are depleted for induced and enriched for spontaneous SVs. ** p-value < 0.01, * p-value < 0.05, post-hoc Fisher’s exact test. g) Induced DSBs occur significantly closer to H3K4me3 (*n* = 122,258), H3K27ac (*n* = 103,673), RNA pol II (*n* = 59,021) and CTCF (*n* = 43,285) peaks compared to the permuted ones (*** p-value < 1e-8, t-test). Spontaneous DSBs show closer proximity to heterochromatin compared to both the induced and permuted sets (* p-value < 0.05, t-test). Box plots show the median, first and third quartiles, and outliers are shown if outside the 1.5× interquartile range.

Since the frequency of interactions within compartments A and B is known to be high (Lieberman-Aiden et al. 2009), we hypothesized that pre-existing long-range contacts between distant compartments would increase the probability of SV formation between these compartments. To this end, we investigated the chromatin compartment on both ends of the SVs in the daughter cells and quantified the proportion of SVs that occurred within (|A-to-A| or |B-to-B|) and between (|A-to-B|) compartments. To compute a background expectation set, we generated a set of random shuffled SVs in the mappable genome, keeping SV size and class and chromosome fixed. Compared to our background model, within compartment SVs are strongly enriched (Figure 2c, right). We found more than half of all TOP2-induced SVs to occur in |A-to-A| compartments (p = 0.0002 compared to random set, post-hoc Fisher’s exact test) and fewer than 10% in |B-to-B| compartments. Spontaneous SVs displayed a more uniform distribution with a relatively higher proportion of within than between compartment SVs, although this did not reach statistical significance when compared to the random background set, potentially due to the relatively low number of spontaneous SVs in our set.

The genomic folding principles can be reconciled at the level of topology-associated domains (TADs), which are genomic loci with a high within contact frequency. TADs are thought to play an important role in controlling and restricting gene regulation. Given the enrichment of SVs occurring between similar compartments, we next explored to what extent these SVs were found to disrupt TADs. In comparison to an expected background distribution, we found a significant enrichment of induced SVs associated with TAD disruption (p = 0.009) (Figure 2d, top). In contrast, spontaneous SVs were depleted for SVs expected to disrupt TADs (p = 0.012), suggesting that SVs associated with TOP2-linked SVs are particularly prone to cause gene regulation changes. TAD boundaries are associated with chromatin loops mediated by the structural proteins CTCF and Cohesin (Hadjur et al. 2009; Sanborn et al. 2015). The loop anchor can be observed on Hi-C maps as focal points with high interaction frequency, and prior work has shown that TOP2 is enriched near these loop anchors (Canela et al. 2017). Consistently, we discovered not only TOP2-induced but also spontaneous DSBs to occur significantly closer to loop domain boundaries and anchors (Figure 2d middle and bottom panel).

These results suggest that our model system can capture many of the mechanistic properties that govern SV formation. Rearrangements in cancer genomes, in contrast, are highly shaped by selective pressure occurring over years to decades, that favor the presence of SVs associated with selection. To examine the association between chromatin conformation and SVs identified in cancer genomes, we analyzed SVs from almost 2,700 tumor samples across 30 different tumor types (Li et al. 2020; ICGC/TCGA Pan-Cancer Analysis of Whole Genomes Consortium 2020). We also undertook separate analysis of 3 common cancer types with high SV burden, namely breast, prostate and uterine cancer. In these cancer genomes, we found overall a more even distribution of SVs occurring in A- and B-compartments (Figure S5c). In agreement with the findings from our model system, a significant majority of SVs occurred within compartments (|A-to-A| and |B-to-B|, p < 0.01 for all cancer types compared to our background model, Fisher’s exact text). Uterine cancer showed a striking enrichment of |A-to-A| SVs, the highest proportion across all 30 tumor types and 24% higher compared to pan-cancer SVs and 60% higher compared to our background model (p < 1e-16, Fisher’s exact test), suggesting that influence of chromatin organization varies across tumor types and that SVs in uterine cancers are particularly associated with contact frequencies between A compartments, associated with open chromatin.

Faulty DNA replication is thought to underlie many complex SVs in cancer genomes through e.g. replication fork-stalling and template switching or replication fork collapse (Quinlan et al. 2010; Canela et al. 2017; Yang et al. 2013). Replication timing (RT) is also strongly linked with genome architecture and contact domains. Early replicating regions are preferentially localized in the interior of the nucleus, in close proximity and often overlapping with A-compartments, whereas late-replicating regions localize to the nuclear periphery and coincide with B-compartments (Ryba et al. 2010; Pope et al. 2014). We performed Repli-seq to explore the influence of RT on SV formation and the changes in chromatin conformation. In agreement with earlier findings, we find a robust correlation between early RT (E-RT, defined as the first quartile of Weighted Average, (WA) values) and A-compartments and between late RT (L-RT, defined as the third quartile of WA values) and B-compartments (Figure S5a-b, Pearson correlation 0.77, p<2.2e-16). TOP2-linked SVs were strongly linked to E-RT and depleted for SVs in L-RT. This was also reflected when analyzing both ends of an SV, where induced SVs were more prominently linked to juxtapositions between two E-RT domains and depleted for juxtapositions between L-RTs (Figure 2e-f). DSBs associated with spontaneous SV formation displayed a more bipartite pattern. Surprisingly, their juxtaposition and SV formation were linked to within RT juxtapositions for both early and late RT, suggesting that spontaneous formed SVs are highly influenced by RT per se and to a lesser extent whether it is early or late RT. We performed a parallel analysis of SVs in cancer genomes, which displayed a distribution similar to the spontaneous SVs with significantly higher within-RT juxtaposition frequency compared to our background model (p < 0.01, Fisher’s exact test). In addition to the enrichment of uterine cancer within compartment juxtapositions, we also observed a particularly pronounced enrichment of SVs associated with juxtaposition between E-RTs in this tumor type, 60% higher compared to pan-cancer and conversely 40% depleted for SVs between early and late RTs compared to pan-cancer analysis (Figure S5d-e, p-value < 2.2e-16, Fisher’s exact test).

Active chromatin is associated with specific histone modifications including H3K27ac and H3K4me3 as well as the presence of RNA polymerase II. We asked whether these epigenetic marks were associated with occurrence of SVs following chromothripsis-induction as well as in the spontaneously occurring set. In agreement with our analysis of contact domains and compartments, we found overall that especially the induced DSBs occurred significantly closer to active chromatin marks and CTCF binding sites. The spontaneous DSBs also occur in closer proximity to marks of heterochromatin (p = 0.025 vs induced, p = 0.035 vs shuffled, t-test) (Figure 2g), suggesting that chromatin marks, which are bound by proteins such as transcription factors or structural proteins, are more likely to harbor DSBs in the vicinity (<10-20 Kbp).

### SVs in regulatory regions cause expression dysregulation of nearby genes

Gene regulation is dependent on distant and proximal cis-regulatory elements (CREs) that loop and interact to facilitate transcription factor, mediator complex and Pol II loading to initiate transcription. Our findings of enrichment of SVs intersecting the transcriptionally active A compartments prompted us to investigate the influence of RNA transcription levels on SV formation. We compared mRNA transcription between the maternal cells and two daughter cells, C29 and DCB2, and separated the genes into active and inactive sets based on the level of expression in the maternal cell lines, i.e. before SV occurrence. We found that actively transcribed genes were associated with nearby SVs across C29 and DCB2 with 4-fold increased risk of SV occurrence if the gene is expressed. We further separated the active genes into high and low expression bins (above and below median expression of active genes), which showed that higher expressed genes were more strongly associated with SV occurrence with an increased odds ratios of 1.91 for C29 (p=0.001) and 1.35 for DCB2 (p=0.217, Figure 3a). SVs can affect gene expression by altering CREs, but it has been difficult to ascertain the extent to which SVs affect nearby gene expression in general. By testing the association between breakpoints and nearby gene expression after correcting for copy number effects (see Methods), we find a significant correlation between breakpoint proximity and differential gene expression (Figure 3b, p=0.01, Pearson correlation coefficient = −0.18), with 10% of breakpoint regions associated with differentially expressed genes in cis and 63 % of genes displaying a significant change in gene expression within a distance of 25 kb (Figure 3c, p=0.018). These results collectively demonstrated that SV formation is strongly associated with prior active transcription (Figure 3d).

**Figure 3:**
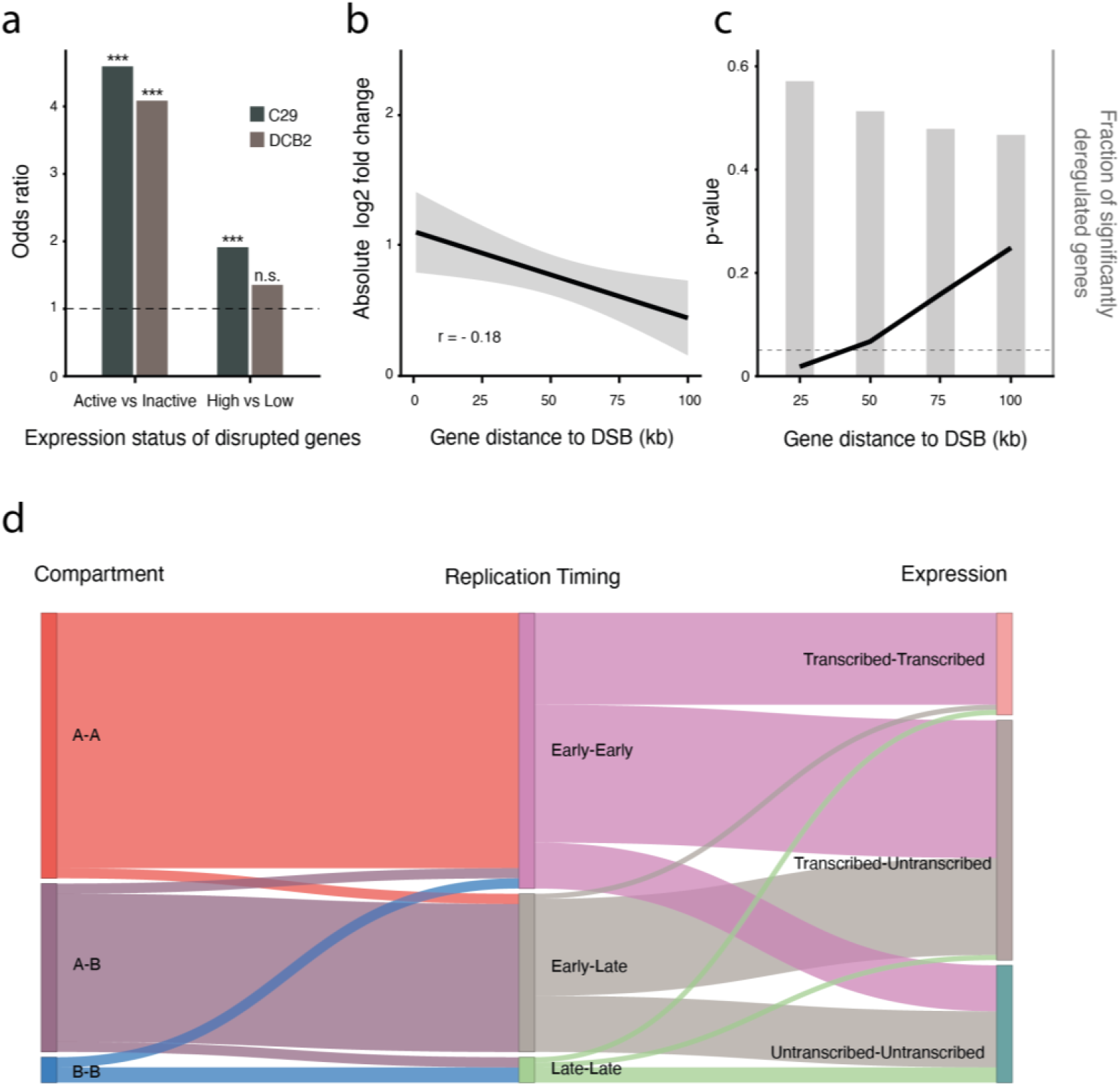
SV-mediated effects on gene expression in *cis*. a) DSBs are associated with highly transcribed genes. (left) Out of 414 DSBs in C29-derived clones, 198 were located within gene bodies; 116 in active, 82 in inactive. The DCB2-derived clone had 137 out of 198 DSBs occur within gene regions; 80 active and 57 inactive. Genes were defined as active if they had TPM > 1 across all replicates. (right) Further subdivision of active genes, links highly (>50%) transcribed genes with the formation of DSBs. ** p-value < 0.01, *** p-value < 1e-5. Fisher’s exact test. b) Absolute log_2_ fold change as a function of distance from the closest DSB cluster in BM175 cells. For every DSB cluster, we considered the gene with the highest absolute log2 fold change in a 10kb sliding window with no overlap. c) P-value estimation of the number of significantly deregulated genes as a function of genomic distance from the most proximal TOP2-linked DSB. Background distribution was estimated by randomly shuffling the gene status (deregulated, non-deregulated. See methods section). Barplots depict the fraction of deregulated genes within the marked genomic window. d) Effects of active chromatin, replication timing and gene expression on the formation of TOP2-linked, induced SVs (*n* = 90). Intermediate replication timing states (*n* = 214) are excluded for visual simplicity. The complete plot is included as a supplementary figure (Figure S6).

### SV-mediated chromatin reorganization affects both gene regulation and DNA replication in *cis*

Genomic compartments disappear during mitosis and reassemble during G1 and S phase (Nagano et al. 2017). Although many RT boundaries coincide with genomic compartments, replication is a dynamic process that occurs during the S-phase of the cell cycle. After having established the importance of compartments and RT in SV formation, we hypothesized that SVs could also impact on the location and type of both features by causing compartment and RT switches. We found several loci associated with such changes in compartment and RT between maternal and daughter cells. A representative example is shown in Figure 4a from the q-arm of chromosome 15 in BM175, which was associated with chromothriptic SVs, including translocations to chromosomes 2, 8, 11 and 12 (Figure S1b-c). Whereas the majority of loci were constant (see, e.g. Figure 4a), we noticed a locus with L-RT and B-compartmentalization in the C29 maternal clone (Figure 4a, red shade), where several DSBs occurred in the BM175 daughter clone (Figure 4a, bottom panel). This was associated with a dramatic switch in replication from late to early RT (measured by delta WA, Figure 4a) and compartment switch from B to A. The SV-associated compartment and RT switch was linked with upregulation of both genes within this locus (WDR72, fold-change=7.5, and UNC13C, fold-change=7.8, Figure 4b) and the emergence of new loop domains (upper right square, shown as open boxes on the Hi-C map, Figure 4b). Most of these new looping contacts coincided directly with sites of DSB, strongly suggesting that they were a direct consequence of the SVs. We found another example of compartment and RT-switch on 15q but in the opposite direction, from A-compartment and E-RT in the maternal C29 clone to a B-compartment and L-RT in BM175 (Figure 4a, green shade). The switch was not associated with DSBs but was accompanied by reduced expression of genes within the compartment and reduced loop contact point strength (Figure 4b, blue shade).

**Figure 4:**
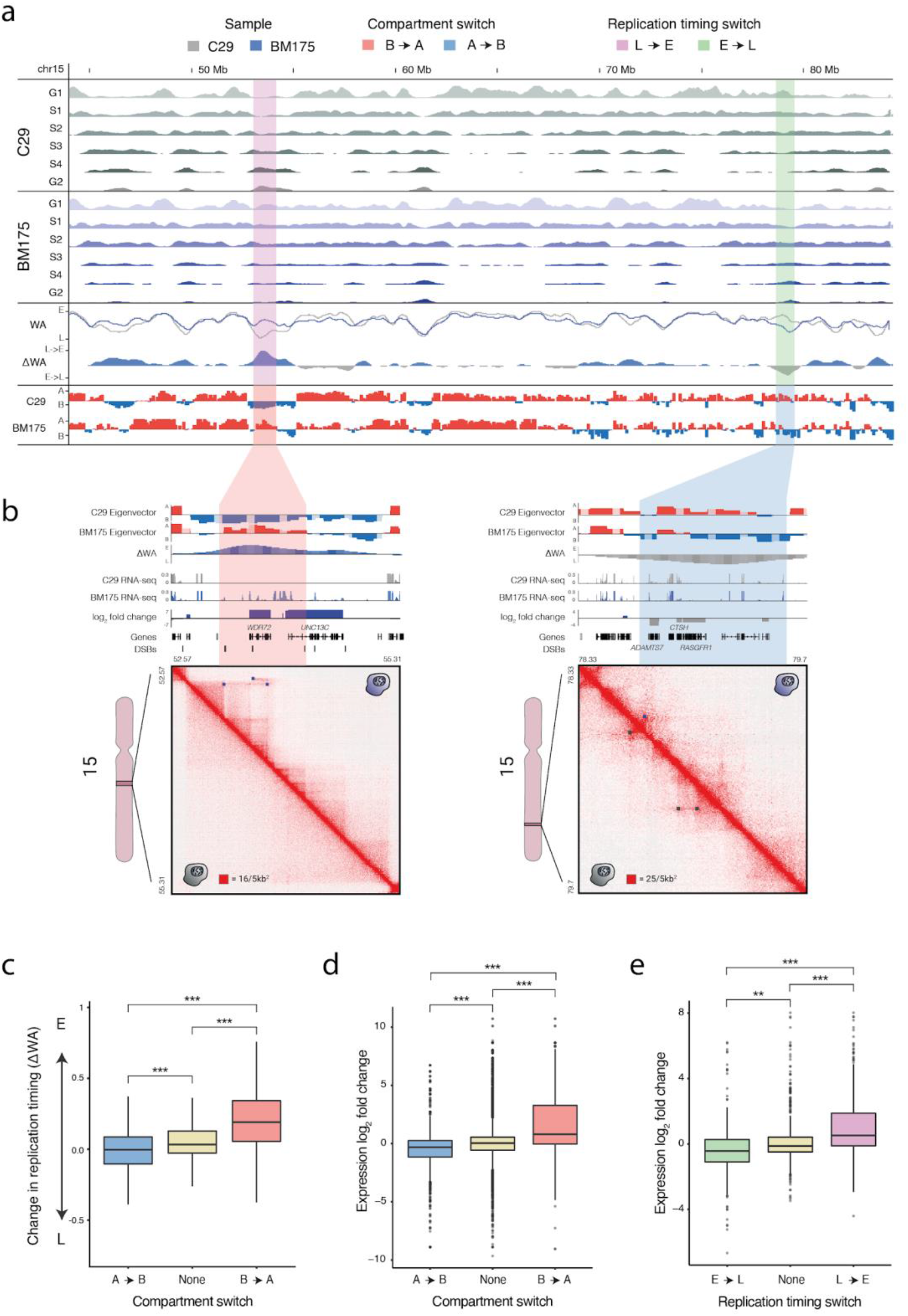
Interplay between compartment switches, replication timing switches and gene expression in BM175 (daughter clone) compared to C29 (maternal clone). a) Representative examples of regions with replication timing switches from Early to Late (green) and Late to Early (pink) coupled with compartment switches from A to B (blue) and B to A (red) respectively. Percent-normalized density values (PNDV) for C29 (grey, maternal) and BM175 (blue, daughter) are shown in the top panels. Weighted average values (WA) and their difference (ΔWa) pinpoint switches in replication timing, similar to the Hi-C eigenvector tracks below for compartment switches. The switches were evenly distributed genome-wide, irrespective of the presence of DSBs. b) Alterations in chromatin structure and expression in regions with a combined compartment and replication switch. (left) B → A and Late → Early switch accompanied with upregulation of WDR72 (log_2_fc = 7.5, p-value = 5e-11) and UNC13C (log_2_fc = 7.8, p-value = 3e-13) and formation of chromatin loops. (right) In contrast, an A → B and Early → Late switch is coupled with downregulation of ADAMTS7 (log_2_fc = −3.9, p-value = 1e-15), CTSH (log_2_fc = −2.2, p-value = 6e-17) and RASGRF1 (log_2_fc = −2.24, p-value = 0.0002), and loss of chromatin looping. c) Genome-wide association of compartment switches and changes in replication timing (ΔWA). A → B (*n* = 298,916) switches have significantly lower ΔWA (E → L) values than regions with no switch (*n* = 1,912,662), and B → A (*n* = 317,686) significantly higher (L → e). *** p-value < 2.2e-16, two-sided t-test. Box plots show the median, first and third quartiles, and outliers are shown if outside the 1.5× interquartile range. d-e) A → B (*n* = 1,798) and E → L (*n* = 327) switches are significantly down-regulating genes compared to regions with no switch, while B → A (*n* = 1,063) and L → E (*n* = 255) upregulate them significantly. ** p-value = 3.9e-7, *** p-value < 7.4e-14, two-sided t-test. Box plots show the median, first and third quartiles, and outliers are shown if outside the 1.5× interquartile range.

To quantify these observations at a genome-wide scale, we computed the extent to which A-to-B and B-to-A compartment domain switches were accompanied by E-to-L and L-to-E RT-switches. In agreement with our observations, A-to-B and B-to-A switches were significantly associated with E-to-L and L-to-E switches, respectively (Figure 4c), both when comparing between the two switch states and when comparing regions without compartment switches (p≪2.2e-16). Having demonstrated the strong preference for SVs to occur in A-compartments and that SVs on average tend to deregulate gene expression upon chromatin domain disruption, we investigated whether compartment switches had a direct impact on gene expression. We found that both A-to-B and E-to-L switches, hypothesized to lead to a more repressed, silent chromatin state, were significantly associated with reduced gene expression. In contrast, B-to-A and L-to-E switches, generally associated with open, active chromatin, were significantly linked to increased gene expression (Figure 4c-d). To distil these findings, we related juxtapositions with compartment, RT and gene activation state, which demonstrates how A-A compartment juxtapositions almost exclusively correlate with E-E RT juxtapositions and that the majority of these SVs involve an active gene at one or both ends (Figure 3d, Figure S6). In contrast, B-B compartment juxtapositions are equally separate between E-E and L-L RT juxtapositions, but nevertheless all involve inactive genes on both ends.

Having established how SVs can change compartments and replication timing to alter gene expression on average, we noticed a striking set of SVs at chromosome 11, which led to novel chromatin looping and a series of dramatic gene expression changes in cis. The locus involved a 90Mb deletion and a 35kb inversion involving BCL9L and CXCR5 (Fig 5). Our Hi-C based derived assembly showed exchange of the regulatory regions between these two genes and additionally a long-range (>900 kb) ectopic loop to BDNF (Figure 5a-c). Integrating allele-specific expression demonstrated simultaneous upregulation of both CXCR5 and BCL9L (>10-fold and 2-fold, respectively, Fig 5d). Conversely, the long-range loop was associated with downregulation of BDNF (2-fold), suggesting emergence of looping with a repressive element or disruption of existing enhancer-promoter interactions at the locus. In short, this locus exemplified how a few SVs can cause altered chromatin organisation and dramatic deregulation of several genes, one of them being BCL9L, a known oncogene (Zatula et al. 2014).

**Figure 5:**
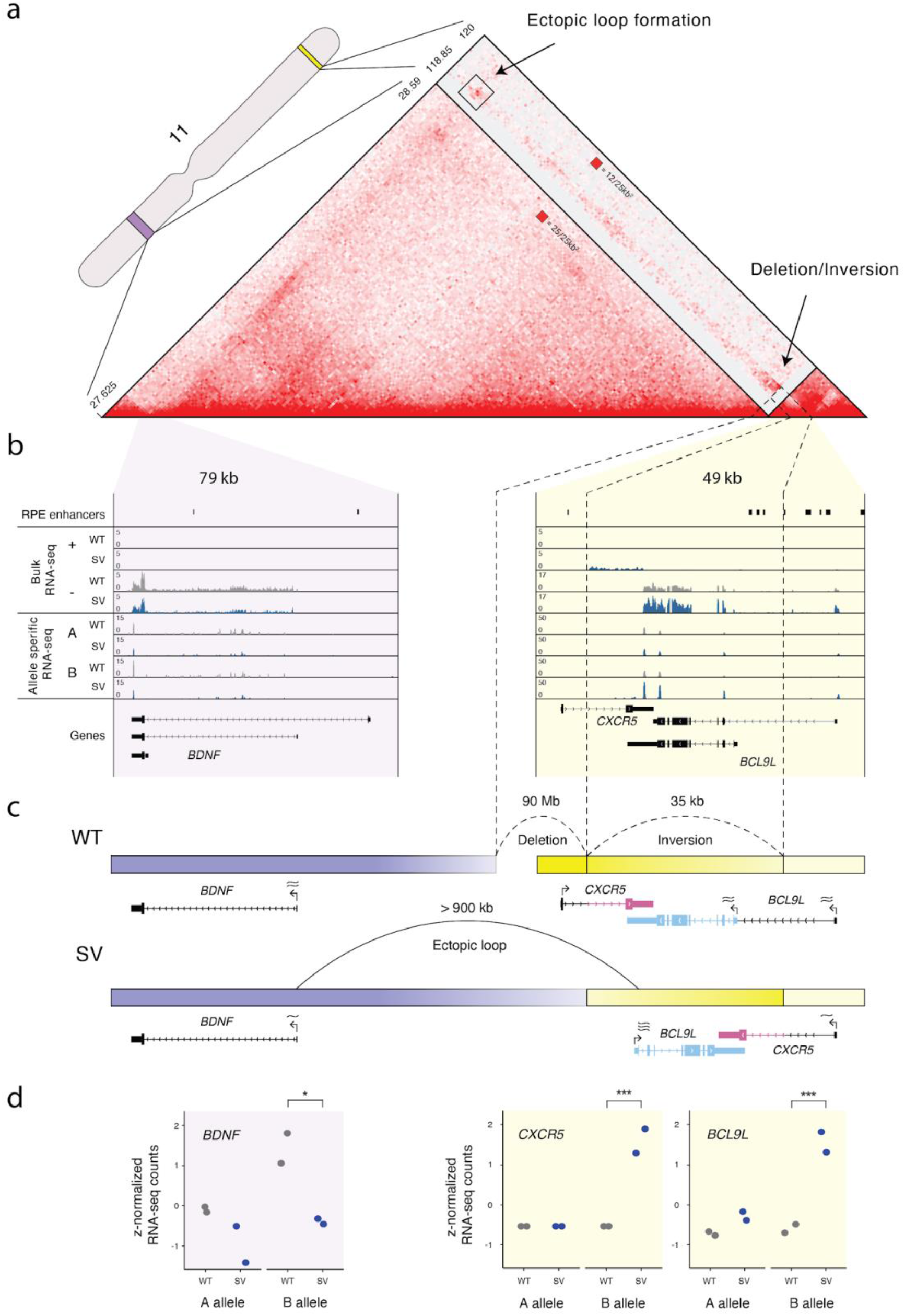
Complex SV form ectopic loops to simultaneously deregulate BDNF, CXCR5 and BCL9L in cis. a) Hi-C map highlighting a complex SV on chromosome 11 in BM175. The SV results in the juxtaposition of the BCL9L/CXCR5 locus (yellow) 90 Mb upstream and the formation of an ectopic loop with BDNF (purple). The genomic regions depicted on the Hi-C maps correspond to the highlighted (purple and yellow) regions on the chromosome ideogram. b) Gene expression tracks stratified by plus/minus strand (bulk RNA-seq) and A/B allele (Allele-specific RNA-seq). SVs are affecting the B allele only. Dashed vertical lines track the breakpoint positions. c) Schematic representation of the resolved assembly and the SV mediated deregulation of BDNF, BCL9L and CXCR5 in cis, as inferred by Hi-C and RNA-seq data. d) Allele-specific gene expression of BDNF, CXCR5 and BCL9L. * p-value < 0.05, *** p-value < 0.001.

In summary, our integrative results using derived assemblies integrated with chromatin organization and replication information implicate a direct and quantitative functional consequence of SVs on genomic architecture, where SVs not only change the chromatin structure but can also have direct consequences on replication and nearby gene expression.

## Discussion

Genomic instability fuels cancer development and can cause complex rearrangements, leading to reordering of genomic segments on the derivative chromosomes. To gain mechanistic insight into SV formation and their consequences on gene regulation, we employed an approach that leverages the power of Hi-C to detect complex SVs and to infer the position and orientation of the rearranged genomic fragments in the derivative chromosomes (Figure 6). While previous reports have described the capacity of Hi-C data to identify SVs (Harewood et al. 2017; Dixon et al. 2018), we present in this study a framework to discover, interpret, and assemble complex SVs. Here, we utilise these highly rearranged and manually assembled genomes to identify general principles of SVs on chromatin organisation and gene regulation.

**Figure 6:**
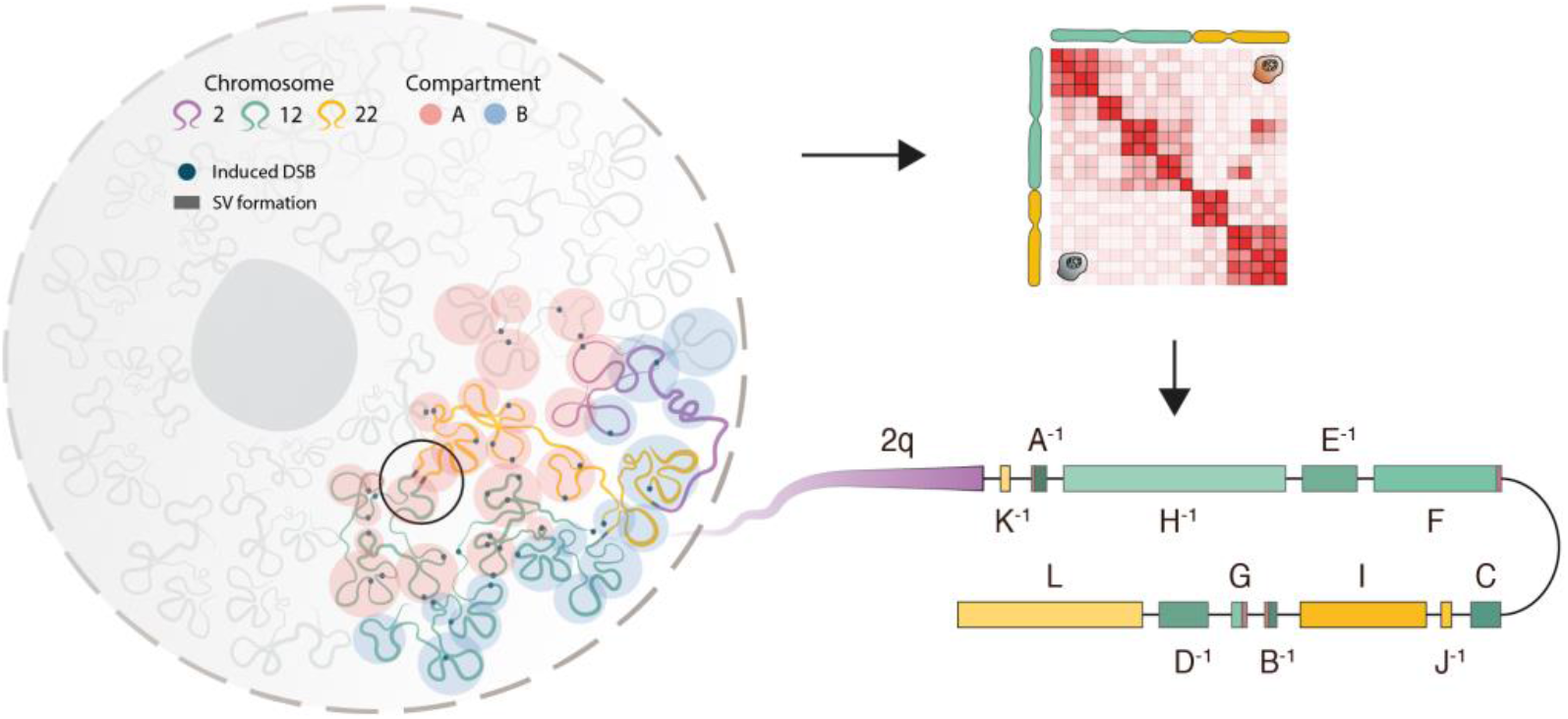
Schematic illustration of genome conformation during DSB formation. The Hi-C map allows us to track SV formation and assemble the derivative chromosomes, which enabled us to study how SVs affect chromatin organisation and gene expression.

The folding architecture of the genome is an important determinant for SV formation, where cell-, histology-type or treatment differences may be at play in shaping differential rearrangement landscapes across different tumor types. As a consequence of the SV formation, significant changes in genome architecture can include chromatin domain disruption, compartment and RT switch with consequences on gene regulation. The 3D genome impacts directly on SV formation which again influences the 3D genome organization and gene expression in cis. Such ‘action-reaction’ effects between chromatin organization and SVs imply that studies of disease-associated rearrangements should be evaluated in the context of the 3D genome architecture.

In this study we used a TOP2 inhibitor doxorubicin as a means to induce SVs and provided evidence that the TOP2-linked SVs are strongly associated with both chromatin architecture, replication timing and transcription. Transcription activity does not correlate with TOP2 activity and CTCF-mediated loop domains generally do not require active transcription (Vian et al. 2018; Canela et al. 2017), suggesting that these two features are additive in driving SV formation. The spontaneous SVs, formed after the induced SVs, displayed a similar though less pronounced pattern, with a noticeable preference for intra-compartment SVs (|A-A| or |B-B|), suggesting that compartment proximity is more important than the identity of the compartment. This was consistent with RT, where SVs were more likely to occur within early or late replicating regions.

To provide further support for our findings from our cell line model, we additionally undertook an analysis of cancer samples across 30 different tumor types to evaluate our findings from the model system in the context of samples that are expected to have undergone many series of rearrangements over time. We found a consistent pattern of higher A-A compartment SVs across the tumor types. Uterine cancers displayed an extreme pattern with high A-A and in particular high levels of E-E RT SVs (60% higher abundance in Uterine cancer compared to our pan-cancer analysis). The nature of this preference for early RT and A-compartments in uterine cancers is unclear, but it is tempting to speculate that mutations in factors involved in replication timing could play a role in shaping this pattern. Indeed, uterine cancers are associated with frequent mutations in RT components, such as RIF1 (Mei et al. 2017) and POLE (Hussein et al. 2015). We note that the uterine tumors are all treatment naive, suggesting that the effects are endogenous to the tumor type and not due to specific drugs(Wadler et al. 2003; Brave et al. 2006; Benson and Miah 2017).

We find a strong influence of active transcription on DSB formation and locus partner selection, and this process could involve key factors in the enhancer-promoter interaction such as transcription factors and bromodomain proteins, Mediator as well as RNA pol II. Rearrangements of active chromatin can lead to gene expression dysregulation, exemplified by a set of complex SVs on chromosome 11, which led to de novo loop formation and simultaneous dysregulation of three genes, one of which is the proto-oncogene BCL9L, a gene known to be upregulated across different cancers and associated with increased proliferation (Mani et al. 2009; Toya et al. 2007; Zatula et al. 2014).

A previous study using the balancer chromosome in D. Melanogaster revealed a small but significant proportion of SVs to affect nearby gene expression (Ghavi-Helm et al. 2019). Here we extend these observations to mammalian cells, and show a significant distance-relationship, with 63% of genes in close proximity of breakpoints to be differentially expressed. TADs have been implicated to play an important role in regulating gene expression and TAD-disrupting SVs can lead to dysregulation of developmental genes (Nagano et al. 2017; Lupiáñez et al. 2015) and activation of oncogenes (Weischenfeldt et al. 2017). Further elucidation of TAD disruption at the Sox9–Kcnj2 locus in mice has implied that TAD-disruption per se does not cause gross changes in gene expression but rather the reorganization of promoters and enhancers through, e.g. inversions (Despang et al. 2019). We find that approximately half of all SVs occur in regions of the genome associated with A compartments, active transcription and early replication and that SVs are highly enriched at loop anchors and sites bound by chromatin-associated factors. The propensity has direct consequence on gene regulation, with a siginficant impact on gene expression in cis. We therefore propose a second determinant for assessing the function of SVs, whereby prior 3D proximity, gene expression level and compartment status of the juxtaposed loci shape the extent to which SVs impact on gene expression at the fused locus. The molecular factors involved in genomic proximity-mediated SV formation, as well as the subsequent consequences on the genome architecture, will need further studies including e.g. depletion and mutation studies. Cohesin, CTCF and WAPL are important factors in generating the genome topology by, e.g. loop extrusion in mammalian cells (Sanborn et al. 2015; Haarhuis et al. 2017; Rao et al. 2017) and perturbing these could provide insights into their relevance in SV formation.

## Methods

### Cell lines

hTERT RPE-1 cells were grown in DMEM/F-12 (Thermo Fischer Scientific, 11320074) medium supplemented with 10% FBS (Thermo Fischer Scientific, 10500064) and Antibiotic-Antimycotic (Thermo Fischer Scientific, 15240062). All cell lines were maintained in appropriate densities and were incubated in a humidity-controlled environment (37°C, 5% CO2). All cell lines tested negative for mycoplasma contamination.

### Preparation of WGS, Mate-pair and RNA-seq libraries

Sequenced reads from WGS and mate-pair were produced in the original study of induced chromothripsis in RPE-1 cells (Mardin et al. 2015). RNA-seq assay for DCB2 was also produced in the original study. For a detailed description, refer to the Materials and Methods section of that paper.

### Preparation of ssRNA-Seq libraries

Total RNA was purified in duplicate from 1 million frozen cells using the Direct-zol RNA MiniPrep kit from Zymo Research (#R2050), following the manufacturer instructions. RNA purity, A260/A280 and A260/A230 were assessed with Nano Drop 2000. RNA integrity was evaluated by electrophoresis on a denaturing agarose gel using 350 ng RNA. Clear 28S and 28S bands were observed indicating no RNA degradation. Before strand specific RNA-seq library preparation in duplicate, using the Directional RNA Library Prep kit for Illumina (NEB#7420), in combination with NEBNext Multiplex Oligos for Illumina Set 1 (NEB#E7335), ribosomal RNA depletion was performed using 1 μg RNA using NEBNext rRNA Depletion kit (NEB#6310). Concentration of final libraries was measured using Qubit 2.0 Fluorometer in combination with Qubit dsDNA HS Assay Kit (Invitrogen). Size distribution and fragment length were assessed using Agilent Bioanalyzer HS-DNA Chips. The molarity of individual libraries was calculated considering the concentrations and average fragment size. Pair-end sequencing of 1.6 pM pooled libraries were sequenced with Illumina NextSeq 500 Sequencing System using NextSeq 500/550 Mid Output v2 kit (300 cycles). PhiX Control (Illumina) was added at 1% to the pool as an internal control before sequencing.

### Preparation of CHIP-Seq libraries

CHIP-Seq libraries were prepared using the CHIP-IT protocol from Active Motif according to manufacturer’s instructions. H3K4me3 antibodies were obtained from Active Motif. Libraries were prepared and sequenced on the Illumina HiSeq 2000 platform with 51 cycles according to the manufacturer’s instructions.

### Preparation of Repli-Seq libraries

Repli-Seq libraries were prepared according to (Hansen et al. 2010) based on a previously described STS replication assay (Hansen et al. 1993). Briefly, 20 million cells were grown to exponential phase and pulsed with BrdU to label the newly replicating DNA. Cells were collected and the cell pellets were resuspended in DAPI staining solution (1 X PBS with 0.1% Triton X-100 and 2μg/mL DAPI). Nuclei were then sorted in 6 fractions (G1, S1, S2, S3, S4 and G2) based on the DNA amount. Sorting was done on a FACS Aria Fusion instrument (BD Biosciences) using a 355 nm laser (450/50 BP). Cells were lysed by the addition of SDS-PK buffer (dH_2_O with 50mM Tris-HCl, pH8.0/ 10mM EDTA/ 1M NaCl/ 0.5%SDS) containing 0.2mg/mL Proteinase K and 0.05mg/mL glycogen for every 100,000 cells collected. The DNA from these cells is isolated by phenol-chloroform extractions and then sonicated into fragments to an average size of ~0.7-0.8 kb using a Covaris S2 instrument (LGC Genomics). Immunoprecipitation was performed using an anti-BrdU antibody (BD Biosciences Pharmigen, Cat.#555627). Precipitated DNA was recovered by another round of phenol-chloroform extraction and then second strand synthesis was done by the random hexamers and exo-Klenow enzyme (NEb). Amount of final DNA was measured by Qubit High sensitivity kit and sequencing libraries were prepared by NEBnext ultra kit (NEb) according to the manufacturer’s instructions.

### Preparation of In-situ Hi-C libraries

All in-situ Hi-C libraries were prepared based on Rao et al., 2014. Briefly, five million RPE cells were crosslinked with 1% formaldehyde for 10 min at room temperature. The crosslinking reaction was quenched with Glycine, then the cells were lysed with Hi-C lysis buffer (10mM Tris-HCl pH8.0 10mM NaCl, 0.2% NP-40) with 1X of protease inhibitors (Sigma complete protease inhibitor cocktail) and the nuclei were recovered. DNA was digested with 100 units of MboI, and the ends of restriction fragments were labeled using biotinylated nucleotides and ligated. After reversal of crosslinks, ligated DNA was purified and sheared to an average length of 400bp using a Covaris S2 instrument (LGC Genomics). DNA was recovered, and the ligation junctions were pulled down with streptavidin beads and prepped for Illumina sequencing using NEBUltra kit.

### Genome reference

Unless stated otherwise, the NCBI GRCh38 (hg38) human genome reference was used for the alignment of all sequenced reads and the GENCODE v30 reference annotation for all human gene related assessments (Frankish et al. 2019).

### Hi-C data processing

All Hi-C datasets were processed using Juicer (Durand et al. 2016) and visualized with Juicebox (Durand et al. 2016). We used hiccups to call loops in C29 with −m 2048 −r 5000,10000 −k KR −f .1,.1 −p 4,2 −i 7,5 −t 0.02,1.5,1.75,2 −d 20000,20000 --ignore_sparsity parameters which produced a set of 8,730 loops at 5kb and 10kb resolution. We performed domain calling on C29 Hi-C maps using arrowhead at 5kb resolution with default parameters and identified 6,163 contact domains with 170kb median size. Pearson correlation eigenvectors were computed by the *eigenvector* module on Knight-Ruiz balanced matrices at 100kb resolution. For every chromosome, the eigenvector sign with the highest association with gene expression and H3K4me3 and H3K27ac, was assigned to the A compartment.

### Allele specific maps

RPE-1 phased SNPs were obtained from a recent study of RPE-1 cell haplotyping from long-range sequencing (Tourdot and Zhang 2019). We examined Hi-C read-pairs with MAPQ ≥ 10 where at least one read was overlapping one or more informative SNP and annotated each informative position to the A or B allele according to the phased information. Reads with varying allele assignments were discarded. Intrachromosomal read-pairs having only one of the reads phased were inferred to the same allele as the phased read. For interchromosomal, we required from both reads to span at least one informative SNP each. Applying these filters yielded allele-specific Hi-C maps with ~12% of the full set of valid Hi-C interactions.

### Allele-specific assembly

Allele-specific merged_nodups.txt files were processed with the 3D-DNA pipeline (Dudchenko et al. 2017) to lift chromosomal coordinates to assembly ones and produce Hi-C maps compatible with Juicebox Assembly Tools (JBAT) (Dudchenko et al. 2018), using the hg38 genome reference as the draft genome.

### Compartment conservation

We used the same process as with the RPE-1 cells to obtain A/B compartment annotations from 8 human cell lines (Rao et al. 2014). Compartment conservation was measured by the concordance of the compartment annotation across the cell lines in 100kb bins. Bins with < 5 compartment concordance were annotated as non-conserved (Nc). Conservation entropy S was calculated as in (Xiong and Ma 2018), and given by the formula:

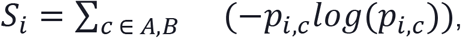

where *p_i,c_* is the fraction of cell lines with annotation *c* for a given 100kb region *i*.

### Copy Number Alterations and ploidy estimates

WGS data were aligned with BWA-MEM v0.7.15 (Li and Durbin 2009). Copy Number Alterations (CNAs) in BM175 and BM178 were identified by Sequenza v3.0.0 (Favero et al. 2015), using WGS from the wild-type C93 as normal. Regions with low mappability score were removed from the analysis. Ploidy estimates were selected based on the Scaled Log Posterior Probability (SLPP).

### Structural Variant calling

#### Mate-pair

Sequenced reads were aligned to the hg19 genome reference with BWA-MEM v0.7.15 and with −T 0 −M parameters. Structural variants were detected by DELLY2 v0.7.7 (Rausch et al. 2012b), using the bam files of the daughter clones as tumor and the matching maternal clones as normal. A high stringency filter was applied to remove SVs detected in ≥ 1% a set of germline SV calls from 1105 healthy individuals from the 1000 Genomes Project (The 1000 Genomes Project Consortium 2015). Furthermore, we required at least 4 supporting reads with MAPQ ≥ 20. High-quality SVs coordinates were converted to the hg38 genome assembly by liftOver (Hinrichs 2006). Complex SVs were identified as described in (Li et al. 2020) and chromothripsis was identified using Shatterseek with default parameters (Cortés-Ciriano et al. 2020).

#### Hi-C

SVs were annotated on the Hi-C map at 5kb resolution in regions of ectopic contacts with high interaction frequency. The exact breakpoint position was identified as the pixel with the highest intensity, located at the origin of the interaction shadow. Ties within the same rearranged fragment were resolved by choosing the position with the highest contact frequency, requiring at least 5 supporting read-pairs with MAPQ ≥ 30 in the given 5×5kb pixel.

The derivative assembly of BM178 was produced using SV calls identified exclusively by Hi-C. For the analysis of DSB and juxtaposition mechanisms, we used SV calls identified by both mate-pair and Hi-C data, except for spontaneous SVs which were only present in the Hi-C maps.

#### PCAWG data

SV calls from 2,693 tumor samples were generated by consensus calling from four SV calling pipelines (ICGC/TCGA Pan-Cancer Analysis of Whole Genomes Consortium 2020; Rheinbay et al. 2020).

### Replication timing

Repli-Seq reads were processed according to Hansen et al. (Hansen et al. 2010) guidelines. The sequenced tag density of each cell-cycle fraction was calculated in 50kb sliding windows at 1kb intervals and normalized to tags per million, excluding intervals with N’s. To account for sequence, mapping and copy-number variation biases by converting the TPM values to percent-normalized density values (PNDV), representing the percentage of total replication occurring on a given 1kb bin within the examined cell-cycle fraction. PNDV across all six cell-cycle fractions were combined into a single Weighted Average (WA) score using the following ENCODE formula:

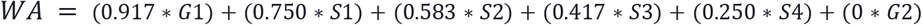

WA reflects Replication Timing (RT), with high values representing early replication and low values late. Regions with WA values lower than the 1st quartile (WA < 0.22) and higher than the 3rd (WA > 0.74) were classified as late RT (L-RT) and early RT (E-RT) respectively.

Repli-seq was performed on C29 on BM175 cells, and WA scores were computed independently for each sample. WA was highly correlated between C29 and BM175 (Pearson correlation 0.81, p ≪ 2e-16). To identify regions with an RT switch, we calculated the difference in replication timing (ΔWa) by subtracting the WA of C29 from BM175 so that positive values represent a transition to Earlier replication timing. We defined regions below the 10% quantile (ΔWA < − 0.11) and above the 90% quantile (ΔWA > 0.29) as E-to-L and L-to-E respectively.

### ChIP-seq

Single-end H3K4me3 ChIP-seq reads generated in this study and publicly available RPE-1 H3K27ac, RNA polymerase II, CTCF, and whole-cell extract (WCe) ChIP-seq reads (GSE60024) we processed by the ENCODE ChIP-seq (https://github.com/ENCODE-DCC/chip-seq-pipeline2), with *--species hg38 --type histone --se*. With the exception of H3K4me3, we used WCE as a control.

### RNA-seq

Single-end and paired-end RNA-seq reads were aligned using *STAR v2.5.3* (Dobin et al. 2013) with --chimSegmentMin 20 and the rest parameters at default. Read coverage was computed at 50bp bins and normalized by Bins Per Million mapped reads with *bamCoverage v3.2.0* from the *deeptools* suite (Ramírez et al. 2016). Transcript expression levels were quantified as Transcripts Per Million (TPM) and collapsed to gene-level expression by *salmon v0.14.0* (Patro et al. 2017) with default parameters and quasi-mapping-based mode. For every gene, we calculated the mean TPM across all available replicates per sample and classified genes with mean TPM < 1 as inactive. Furthermore, we classified active genes as high or low based on the quantile of the TPM distribution of the active genes; high ≥50%, low < 50%. Differential gene expression between technical replicates of BM175 and C29 was performed by the R package *DESeq2 v1.26.0 (Love et al. 2014)*. To minimize effects of copy number alterations we adjusted the gene read counts of BM175 based on the respective gene copy number state as estimated by *sequenza*. Distance effect of DSBs to gene expression was measured as a function of log_2_ fold change of the gene with the highest log_2_ fold change within non-overlapping 10kb bins from a DSB. To avoid biases introduced by genomic loci with multiple breakpoints, we clustered DSBs that resided within a 50kb distance from each other with *bedtools v2.26.0* (Quinlan and Hall 2010). Allele specific gene expression was evaluated as in (Ghavi-Helm et al. 2019). Briefly, RNA-seq reads that overlapped one or more informative RPE-1 SNPs were assigned to the respective haplotype. Reads with ambiguous allele assignment were discarded.

### Cis effects/p-value estimation

To assess the effects of SVs to nearby gene expression we measured the frequency of significantly differentiated genes (q < 0.05) that are proximal to an SV setting the distance cut off to 25, 50, 75 and 100kb. To obtain an empirical p-value, we estimated the background frequency model by randomly shuffling the gene status to differentiated/non-differentiated 10,000 times and evaluated the p value by counting the number of random shuffled genes with an effect larger than the observed.

### Significance testing

Odds ratio scores and p-values were estimated using Fisher’s exact test unless stated otherwise. Empirical distributions of SVs were estimated by shuffling the induced SVs on the same chromosome as the observed set of SVs using *bedtools*, excluding telomeres and centromeres, and preserving the size of intrachromosomal SVs. Each shuffle was performed 1,000 times.

### Visualization

Hi-C maps were produced by *Juicebox v2.0.0* (Durand et al. 2016) and genome browser tracks by the Integrative Genomics Viewer, *igv v2.5.3* (Robinson et al. 2011). All other plots were produced in *R v3.6.0* (R Core Team 2019) using the *base*, *ggplot2* (Wickham 2016), *circlize* (Gu et al. 2014) and *flipPlots* packages.

---

**Figure S1:**
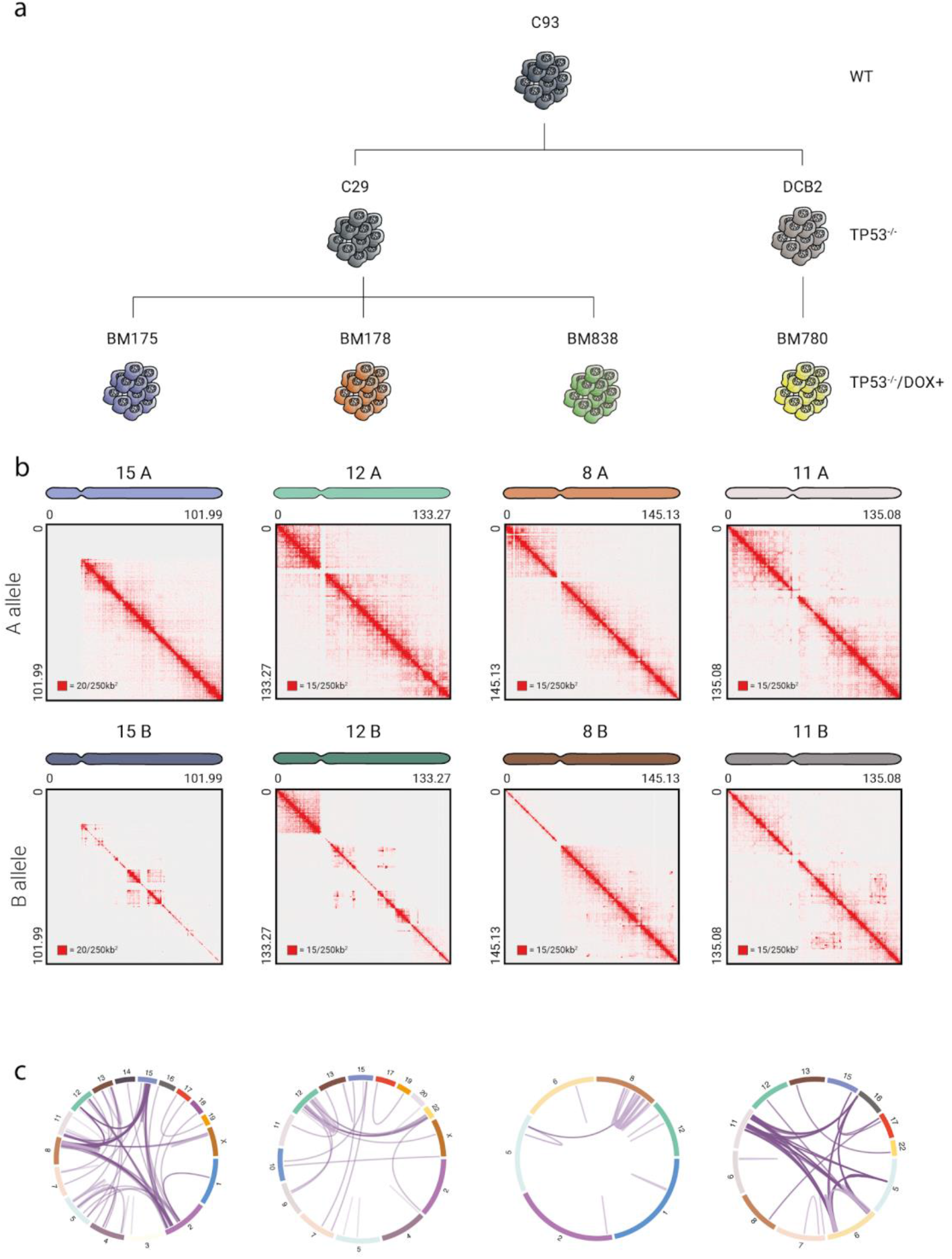
Human cell-line model system Hi-C analyses. a) A 3-tier model system to study SVs. C93 cells (WT) were used to generate two populations of *TP53*-deficient cells, C29 and DCB2 (*TP53*^−/−^). Upon DSB induction with doxorubicin on *TP53*-deficient cells, we generated four distinct populations carrying complex genomic rearrangements (*TP53*^−/−^DOX+), BM175, BM178 and BM838 derived from C29 and BM780 from DCB2. b) Allele-specific Hi-C maps of whole chromosomes carrying complex rearrangements, corresponding to the TP53^−/−^ DOX+ samples in the panel above. c) Genome-wide SVs of the *TP53*^−/−^DOX+ samples, BM175 (*n* = 138), BM178 (*n* = 44), BM838 (n = 25) and BM780 (*n* = 99). The set of SVs used is a union of SV calls from whole-genome sequencing data and Hi-C.

**Figure S2:**
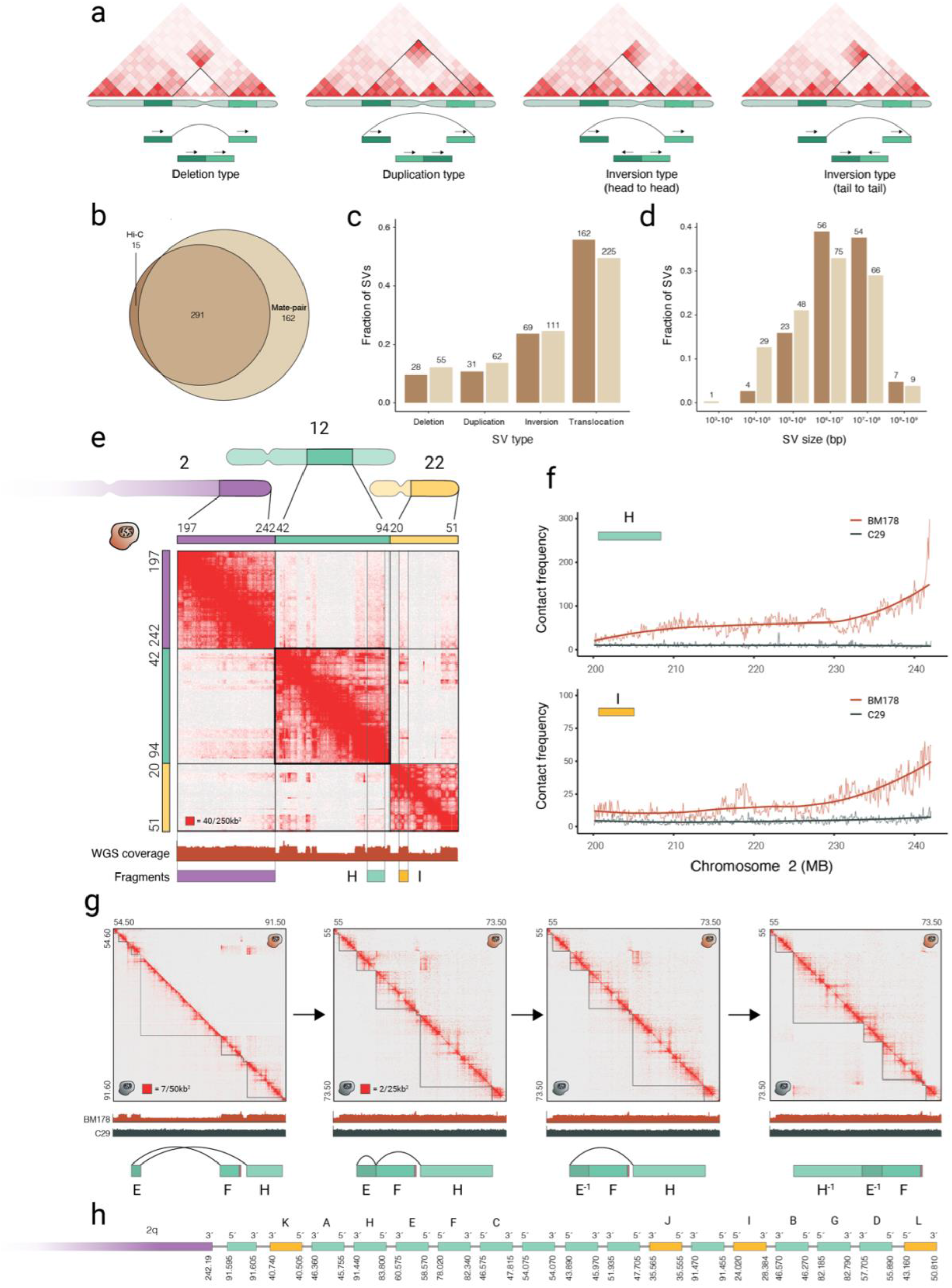
Hi-C-based SV calling and assembly. a) Footprints of deletion-, duplication- and inversion-type SVs on a Hi-C map. b) Overlap of Hi-C and Mate-pair SV calls. Mate-pair SV calls were extended 2.5kb upstream and downstream each breakpoint to match the 5kb resolution of Hi-C SV calls. c-d) Fraction of each rearrangement type (c) and size distribution (d) in Hi-C (brown) and Mate-pair (beige) SV calls. e) Partial map of chromosomes 2, 12 and 22. A faint signal on the telomere of chromosome 2 provides evidence of direct contact with rearranged fragments from chromosome 12 and 22, annotated H and I respectively. The genomic regions depicted on the Hi-C maps correspond to the highlighted regions on the chromosome ideograms. f) Average contact frequency of fragments H and I with the q arm telomere of chromosome 2 for BM178 and the maternal C29 cell line. g) Assembly of fragments E, F and H using Juicebox Assembly Tools (JBAT). All plots display a partial view of the B allele of chromosome 12 with BM178 on upper and C29 on the lower triangle. The illustrations below the plots follow the fragment position in the respective assembly. In a stepwise procedure, we split the chromosome on the predicted sites (left), and remove the deleted regions (left-center). We invert fragment E (right-center) to resolve the first rearrangement, followed by inversion and repositioning of fragment H (right) to resolve the second and last rearrangement. Ectopic domains form between the fragments in the derivative assembly. h) Derivative 2-12-22 with all segments of chromosome 12 included. Segment sizes are not scaled to sequence length. Genomic coordinate annotations are in megabasepairs.

**Figure S3:**
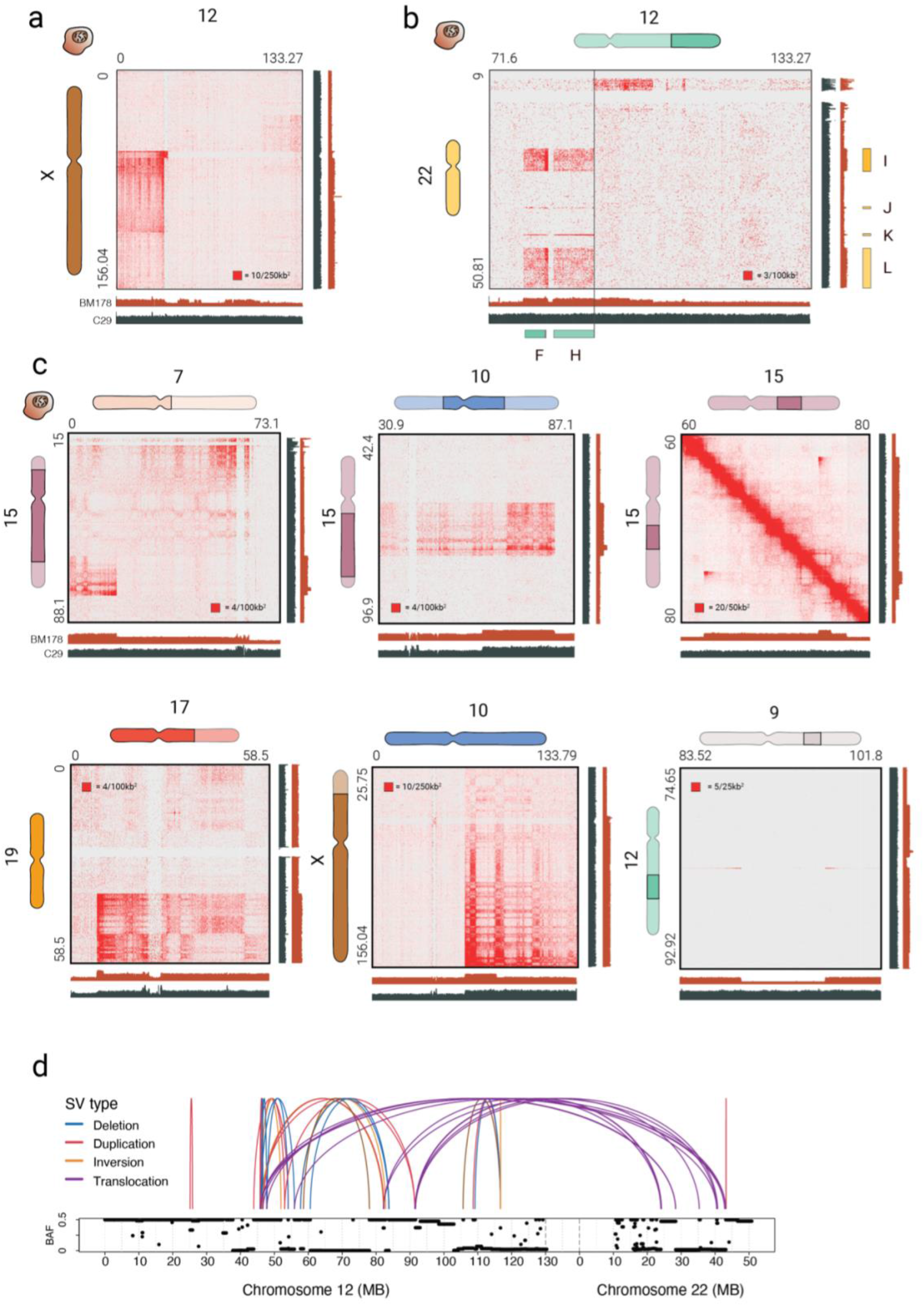
Complex inter- and intra-chromosomal SV detection by Hi-C. a) Centromere joining of 12p and Xq b) Evidence of the remaining 12q residing along the 22p arm. The enriched contacts downstream of fragment H align with changes in copy-number in WGS data. Due to the low complexity of 22p, we were unable to identify interactions at higher resolutions. c) Additional, large scale rearrangements between several chromosomes of BM178. d) Whole-genome view of allele-frequency (top) and read depth ratio (bottom) estimated by Sequenza on WGS data. Arches and colors represent types of SVs within and between chromosome 12 (left) and 22 (right). The genomic regions depicted on the Hi-C maps correspond to the highlighted regions on the chromosome ideograms.

**Figure S4:**
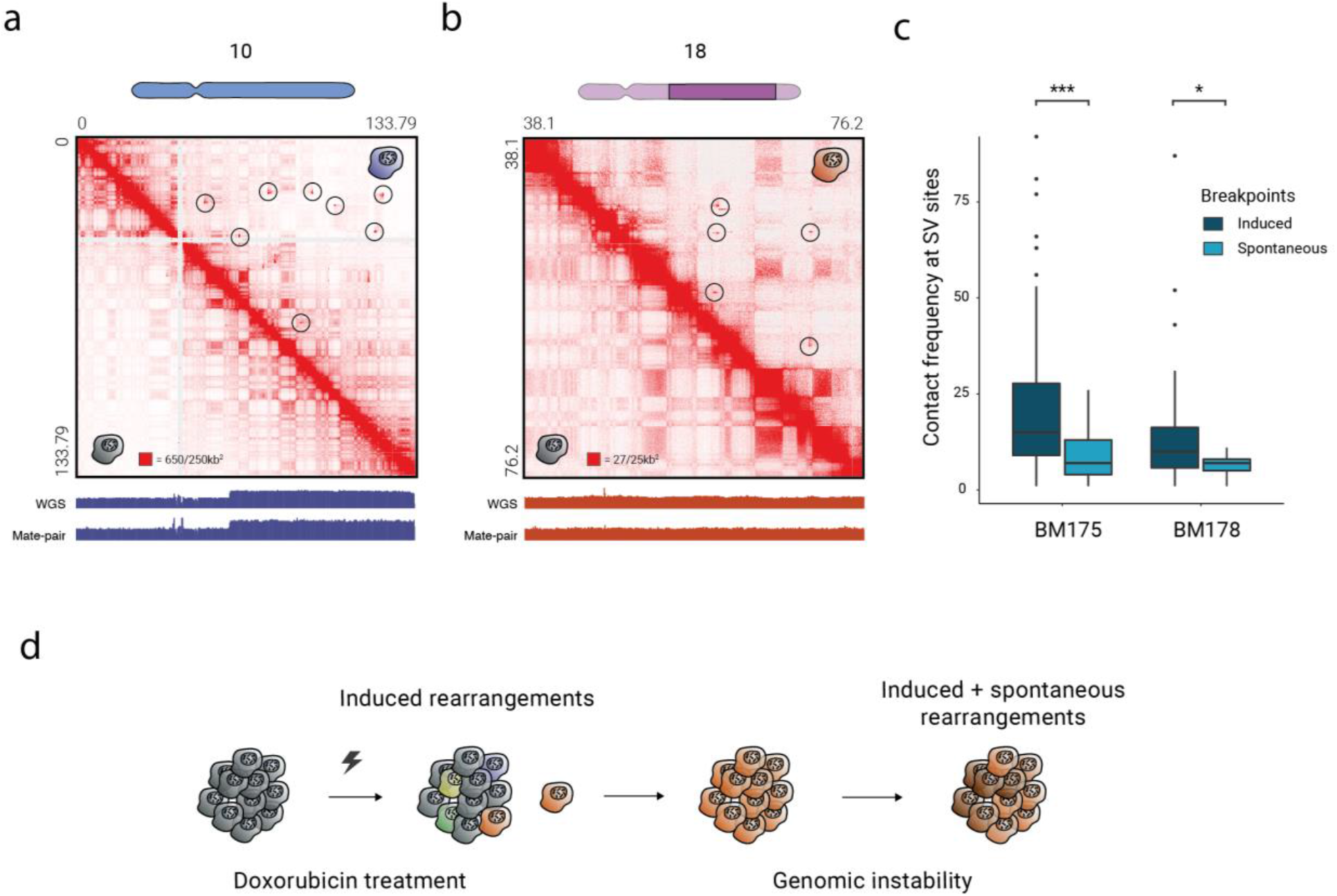
Induced and spontaneous SVs in BM175 and BM178. a-b) Structural variants in BM175 chromosome 10 (a, *n* = 13) and BM178 chromosome 18 (b, *n* = 9) detected in Hi-C data with no evidence on WGS or mate-pair support. The genomic regions depicted on the Hi-C maps correspond to the highlighted regions on the chromosome ideograms. c) Comparison of contact frequency at juxtaposed loci between induced and spontaneous rearrangements. Frequencies are computed at 5kb resolution within a 25×25kb window around the juxtaposed locus. Mann-Whitney U one-sided tests between the two distributions reveal significant contact depletion in the spontaneous events (p-value = 0.003 and 0.03 in BM175 and BM178 respectively). Box plots show the median, first and third quartiles, and outliers are shown if outside the 1.5× interquartile range. d) Formation of DSBs as a result of doxorubicin treatment (induced) and genomic instability (spontaneous).

**Figure S5:**
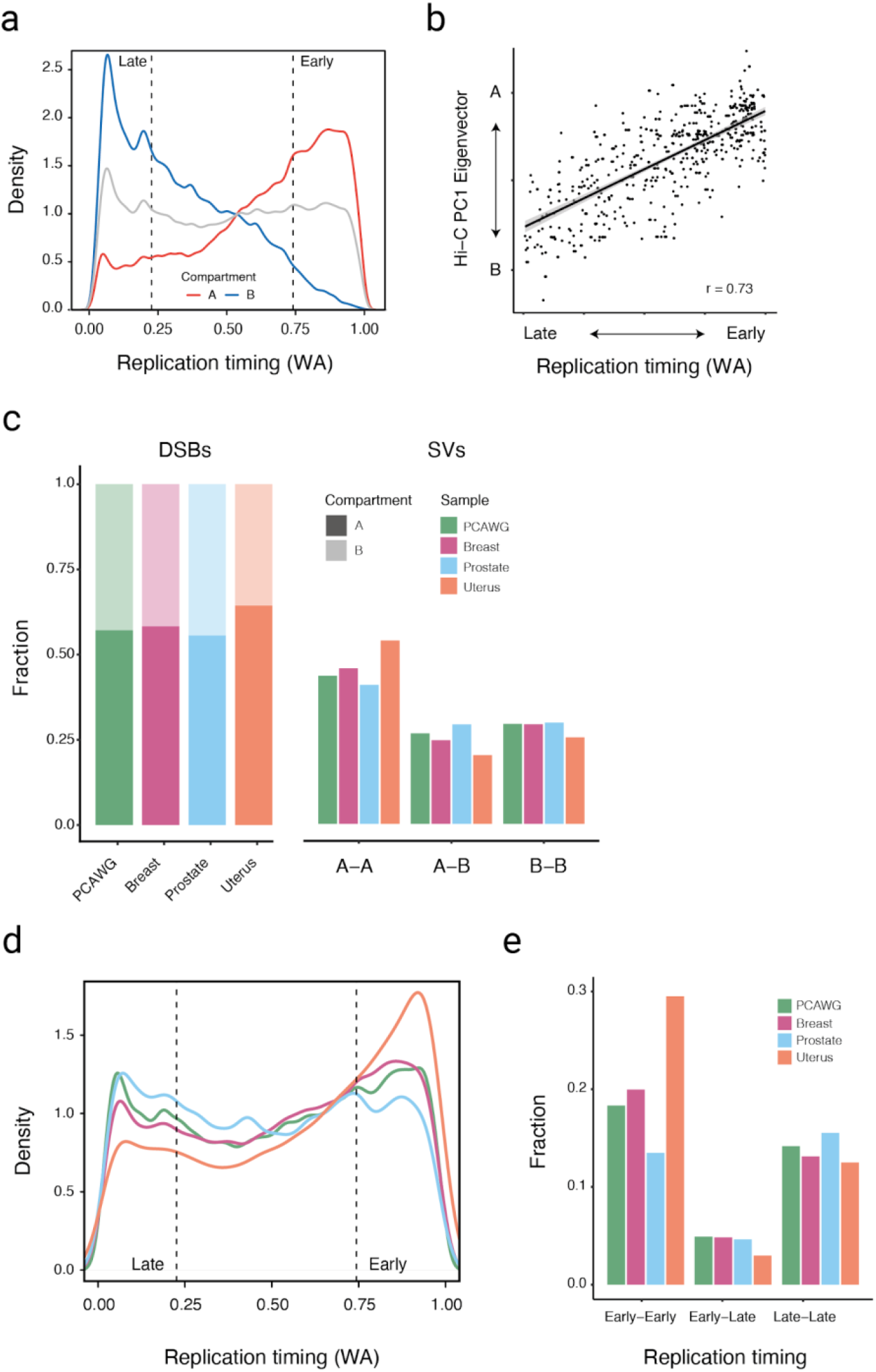
Association of somatic rearrangements in 2,700 tumor samples with chromosomal compartments and replication timing. a) Distribution of replication timing weighted averages (WA) scores in the active A (red) and inactive B (blue) chromosomal compartments. Grey line shows the average WA score genome-wide. Values below the 25% and above the 75% quantile (dashed vertical lines) are assigned to early and late replication respectively. b) A/B compartment scores are highly correlated with replication timing (Pearson correlation = 0.77, p-value < 2.2e-16). Compartment scores are represented by the first principal component eigenvector of the observed/expected Hi-C map. c) Distribution of DSBs (left) and SVs (right) on A and B compartments in pan-, breast, prostate, and uterine cancer samples. Uterine tumors show an enrichment in A-to-A juxtapositions. d,e) Distribution of DSBs (d) and SVs (e) with regards to replication timing. Uterine samples show a striking abundance of Early-to-Early SVs, suggesting an effect of replication timing in SV formation. Early-to-Late SVs are depleted overall.

**Figure S6:**
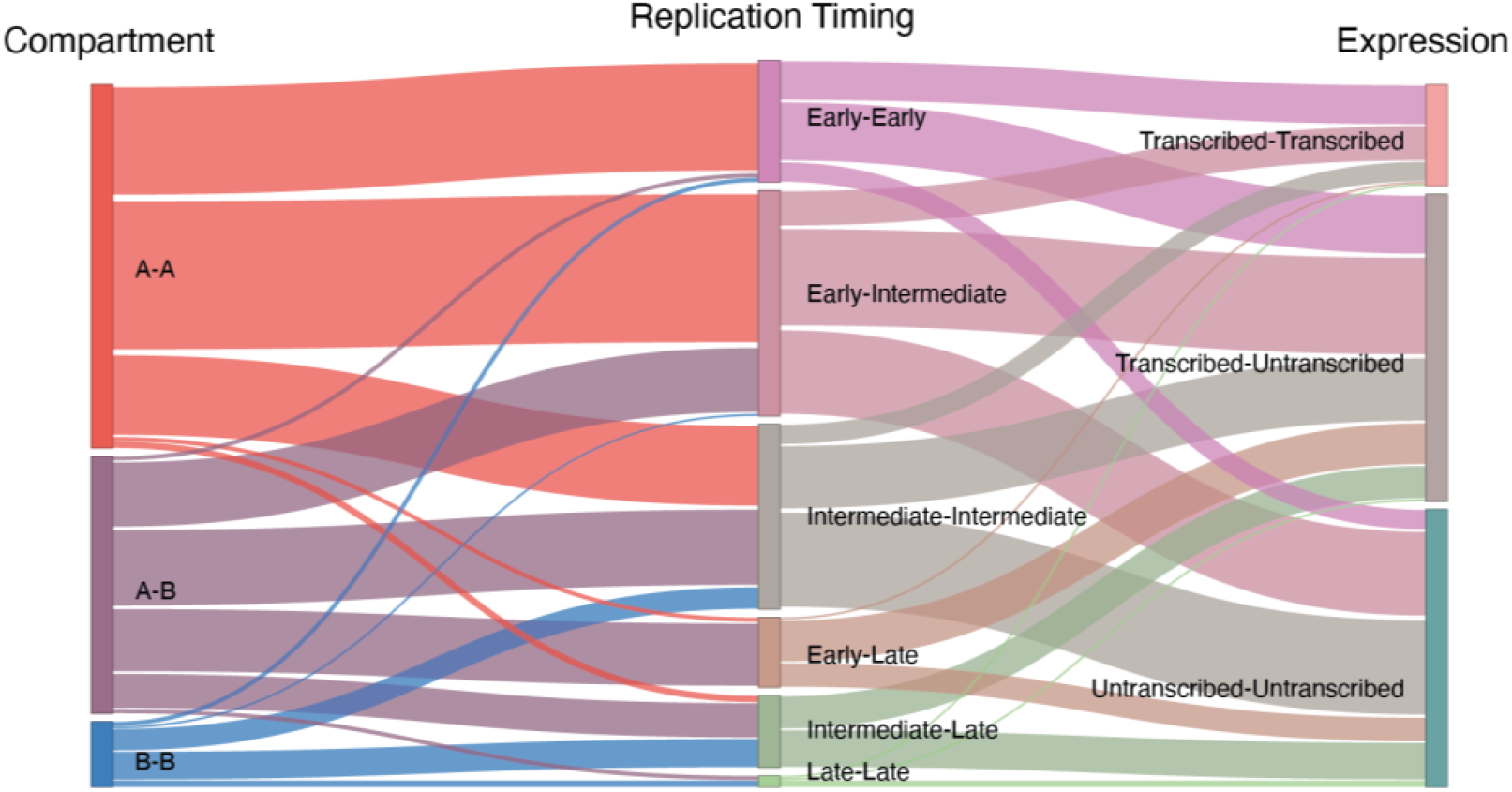
Association between chromatin compartments, replication timing and gene expression on the formation of TOP2-related, induced SVs (*n* = 304). In the gene expression column, ‘Transcribed-Transcribed’ refers to both loci of the SV being associated with transcribed genes, and vice versa.

## Data access

Datasets are available through the European Nucleotide Archive (ENA) under study accession number PRJEB40747. Code used in this study is available at https://bitbucket.org/weischenfeldt/sv_3dchromatin_paper

## Competing interests

The authors declare no competing financial interests.

## Acknowledgements

We thank Ivan Bochkov for helpful comments on the manuscript and Cheng-Zhong Zhang for providing SNP-phased RPE-1 genome data. JW and NS was partly funded by a grant from the Independent Research Fund Denmark (0134-00265b). Members of the Korbel group received funding from the European Research Council (ERC, 336045).

## Author contributions

Conceptualization: JW, NS; Hi-C based assembly Methodology: NS, SH, ELA, JW; RPE-1 cell line generation and perturbation: BM, JOK; Hi-C experiment: BM, FGRG; Hi-C data analysis: NS; ChIP experiment: BM, AS; ChIP data analysis: NS; RNA-seq experiment: BM, FGRG; RNA-seq data analysis: NS; Repli-seq experiment: BM; Repli-seq data analysis: NS; Hi-C based assembly: NS; Integrative data analysis: NS; Visualization: NS; Manuscript, first draft: JW, NS; Manuscript, revisions: JW, NS, BM, AS, JOK; Project Administration: JW; Funding acquisition: JW, JOK; Supervision: JW, ELA

## Notes

### Competing Interest Statement

The authors have declared no competing interest.

